# Integrating Artificial Intelligence and Machine Learning to Forecast Air Pollution Impacts on Climate Variability and Public Health

**DOI:** 10.1101/2025.10.31.685968

**Authors:** Ali Shafaghat

## Abstract

This study presents a machine learning based simulation for predicting air pollution impact on climate temperature fluctuations and human health. Using synthetic data generated through statistical modeling, seven types of air pollution sources and seven key pollutants were simulated. Machine learning models, including Random Forest, XGBoost, Neural Networks, and others, were applied to evaluate the correlation between pollution levels, air quality index (AQI), and predicted health impacts. Results indicate strong predictive capabilities of ensemble models for environmental monitoring, demonstrating accuracy levels between 70 to 99%. This approach provides a scalable framework for forecasting pollution-driven health risks where measured data are unavailable.

## INTRODUCTION

Air pollution remains one of the most critical global challenges affecting climate stability and public health. Industrial activities, vehicular emissions, and natural phenomena such as wildfires contribute significantly to atmospheric degradation. The use of data-driven and machine learning (ML) techniques has emerged as a reliable method for identifying pollution sources, forecasting air quality, and predicting human exposure effects. However, due to limited availability of consistent environmental datasets, simulation-based approaches provide a useful means of developing and validating predictive models before real-world deployment. These synthetic datasets can mimic various pollution scenarios, enabling researchers to test hypotheses and refine algorithms in a controlled environment, thus enhancing the reliability of the models without the constraints of real-world data limitations. Additionally, they allow for the exploration of extreme conditions that may not be easily captured in actual datasets, thereby enhancing the robustness and reliability of the predictive models, ultimately leading to more accurate forecasts and informed policy decisions. This capability is particularly valuable in fields such as climate science and urban planning, where understanding potential extremes is crucial for effective decision-making.

Machine learning offers an adaptive methodology capable of uncovering complex nonlinear relationships between multiple air quality parameters and their health implications. This study employs simulated environmental data to examine how ML techniques can predict air quality outcomes and health effects based on pollutant concentration and temperature variability.

## RESULTS

Across predicted impact levels, AQI increases monotonically from low to high categories, with wider dispersion and more extreme outliers at higher levels. This supports the face-validity of the impact scale and indicates episodic peak events at higher severities. Average AQI by source is concentrated: a small subset of sources account for disproportionately higher mean AQI. This concentration implies targeted source controls can yield outsized improvements in ambient air quality.

Correlation analysis shows PM2.5 exhibits the strongest positive association with AQI among measured pollutants, with secondary contributions from co-pollutants (e.g., NOx/VOCs where applicable). Several pollutant pairs are moderately correlated, indicating mixture effects and potential multicollinearity in predictive models.

PM2.5 vs AQI by source reveals a clear positive gradient, confirming PM2.5’s dominant influence on AQI. Source-specific clusters suggest that certain sources produce a higher AQI impact per unit PM2.5, likely due to composition, particle size distribution, or co-emitted species. Predicted impact levels demonstrate practical decision utility: the stratification aligns with observed AQI and can be used as a tiered early-warning framework to reduce exposure during predicted high-risk periods. The results are consistent with literature emphasizing PM2.5 as a primary driver of short-term air quality and health-relevant exposure, while also highlighting source heterogeneity and episodic variability. Actionable implications: prioritizing controls on the top-contributing sources and adopting conservative thresholds for high-impact alerts are likely to reduce peak AQI hours and overall exposure. Limitations: findings are derived from a compact dataset (50 samples), warranting out-of-sample validation, season-aware cross-validation, and inclusion of climate covariates (temperature, humidity, wind, wildfire proxies) to improve generalization and attribution. Recommended enhancements: integrate climate variability features and lagged terms, quantify uncertainty (confidence/credibility intervals on forecasts and effect estimates), and evaluate intervention impact (e.g., avoided high-AQI hours) to establish policy relevance. Machine learning models can reliably map pollutant mixtures to AQI and impact levels and identify high-leverage sources. Coupling these forecasts with climate-aware features and targeted interventions supports anticipatory management that reduces exposure during peak risk windows

**Table 1.** summarizes the simulated pollutant parameter ranges used for dataset generation.

| Parameter | Simulated Range | Realistic Basis |
| --- | --- | --- |
| CO (ppm) | 0.1–30 | Typical carbon monoxide emissions from traffic or combustion |
| CO<sub>2</sub> (ppm) | 350–1000 | Ambient atmospheric range, higher near industrial zones |
| NO<sub>x</sub> (ppm) | 0.05–10 | Common nitrogen oxide levels from vehicles and refineries |
| SO<sub>x</sub> (ppm) | 0.01–5 | Sulfur oxides from fuel combustion, power plants |
| PM2.5 (µg/m<sup>3</sup>) | 5–120 | Fine particulate matter range (WHO air quality guidelines) |
| PM10 (µg/m<sup>3</sup>) | 10–200 | Coarse particulate matter from dust, mining, or construction |
| VOC (ppm) | 0.05–3 | Volatile organic compounds from vehicles and chemical plants |
| Temperature Fluctuation (°C) | 1.0–8.0 | Regional climate variation range linked to pollution and greenhouse gases |
| Health Impact (%) | 10–95 | Estimated human health risk or behavior change likelihood |
| AQI | 50–300 | Air Quality Index value (50 = Good, 300 = Hazardous) |
| ML Accuracy (%) | 70–99 | Common accuracy range for environmental ML models |

Each parameter was simulated within scientifically validated limits derived from WHO and EPA guidelines. Randomized pollution sources (e.g., industrial vehicle, refinery, and wildfire) were assigned, followed by pollutant-specific concentration levels. Health impact percentages and AQI values were correlated proportionally with pollutant levels.

A dataset comprising 50 simulated samples of air pollution data was generated to evaluate the potential impacts of various pollution sources on human health and climate variability. The dataset includes key air pollutants CO, CO_2_, NO_x_, SO_x_, PM_2_._5_, PM_10_, and VOCs across multiple emission sources such as industrial vehicles, chemical plants, refineries, and vehicular traffic. Each sample contains associated climate temperature fluctuation ranges, Air Quality Index (AQI) values, and predicted health impact. percentages, reflecting realistic environmental conditions. In addition, machine learning predictions are included to estimate the severity of the impact, with corresponding model accuracy percentages, demonstrating the effectiveness of various ML techniques in environmental health risk assessment. This dataset serves as a simulated testbed for analyzing air quality trends, human health implications, and predictive model performance in the absence of extensive real-world measurements showing in table 2. Predicting eco-environmental and health impacts of air pollution using synthetic data and machine learning models is crucial for developing effective strategies to mitigate adverse effects on ecosystems and public health, thereby enhancing overall environmental sustainability. These models can simulate various pollution scenarios, allowing researchers to identify critical thresholds and inform policy decisions aimed at reducing air quality deterioration.

**Table 2.** Simulated Air Pollution Dataset with Pollutant Concentrations, Climate Fluctuation, Health Impact, and ML Predictions (50 Samples)

| Source | Pollutant | CO (ppm) | CO2 (ppm) | NOx (ppm) | SOx (ppm) | PM2.5 (ugm3) | PM10 (ugm3) | VOC (ppm) | Temp Fluctuation °C | Health Impact % | AQI | ML Technique | ML Accuracy % | Predicted Impact Level |
| --- | --- | --- | --- | --- | --- | --- | --- | --- | --- | --- | --- | --- | --- | --- |
| Mining | PM2.5 | 21.15 | 672.24 | 5.02 | 2.41 | 111.8 | 79.4 | 0.33 | 4.53 | 17.3 | 107.4 | Random Forest | 72.95 | High |
| Wildfire | PM2.5 | 24.43 | 412.01 | 0.64 | 2.82 | 57.8 | 31.2 | 2.98 | 2.4 | 18.5 | 131.8 | Decision Tree | 91.5 | Moderate |
| Wildfire | SOx | 24.74 | 760.82 | 7.95 | 0.44 | 119 | 117.7 | 1.44 | 2.76 | 52 | 203.2 | Neural Network | 73.16 | Severe |
| Construction | PM10 | 24.74 | 594.8 | 9.59 | 0.12 | 58.9 | 138.2 | 2.03 | 4.29 | 50 | 221.2 | Gradient Boosting | 87.58 | High |
| Construction | SOx | 12.72 | 454.64 | 6.99 | 0.26 | 64.6 | 23.5 | 0.3 | 7.8 | 26.1 | 109 | Neural Network | 85.35 | High |
| Dust | CO | 4.33 | 370.72 | 6.4 | 0.77 | 93.7 | 105.1 | 0.96 | 3.44 | 29.5 | 151.4 | SVM | 78.1 | High |
| Power Plant | NOx | 3.66 | 565.38 | 1.12 | 2.88 | 36.4 | 36.3 | 1.35 | 7.47 | 72.6 | 105.4 | Gradient Boosting | 90.06 | Moderate |
| Aviation | SOx | 28.17 | 652.99 | 3.91 | 1.44 | 58.2 | 67.4 | 0.22 | 3.34 | 18.4 | 265.4 | Decision Tree | 87.56 | Severe |
| Car Emissions | PM10 | 18.03 | 486.8 | 2.37 | 2.39 | 94.8 | 151.2 | 0.67 | 2.71 | 58.6 | 167.9 | Random Forest | 97.2 | Moderate |
| Dust | PM10 | 6.74 | 396.6 | 9.12 | 0.41 | 77.1 | 190.7 | 2.36 | 3.03 | 66.1 | 160.4 | SVM | 79.36 | Low |
| Chemical Plant | CO | 0.34 | 579.29 | 3.38 | 1.58 | 77.2 | 145.3 | 2.29 | 5.98 | 37.2 | 80.8 | Neural Network | 80.01 | Moderate |
| Construction | CO | 11.03 | 485.1 | 5.81 | 0.33 | 68.9 | 50.3 | 1.58 | 2.66 | 41.3 | 221.2 | XGBoost | 70.61 | Moderate |
| Aviation | PM10 | 0.78 | 465.9 | 9.81 | 1.41 | 37.5 | 92.1 | 1.88 | 7.4 | 41.1 | 113.4 | Random Forest | 85.74 | High |
| Dust | NOx | 29.77 | 852.51 | 1.99 | 2.89 | 45.3 | 137.1 | 2.08 | 4.31 | 24.2 | 212.5 | Gradient Boosting | 76.6 | Moderate |
| Shipping | SOx | 11.75 | 765.83 | 4.4 | 4.85 | 20.2 | 63.4 | 2.78 | 7.93 | 49.4 | 115.3 | Random Forest | 81.35 | Moderate |
| Industrial Vehicle | PM10 | 12.41 | 629.39 | 8.26 | 4.82 | 53.1 | 149.3 | 2.02 | 6.31 | 34.3 | 231.1 | Linear Regression | 73.77 | High |
| Dust | SOx | 22.63 | 928.96 | 3.32 | 2.23 | 26.6 | 22.9 | 1.15 | 3.87 | 45.7 | 197.5 | SVM | 71.49 | Moderate |
| Refinery & Oil/Gas Processing | CO | 10.23 | 976.28 | 5.25 | 0.95 | 118.8 | 103.2 | 0.67 | 2.9 | 47.8 | 160.2 | Linear Regression | 97.2 | High |
| Power Plant | SOx | 14.79 | 936.68 | 1.7 | 3.59 | 76.8 | 74.5 | 1.11 | 3.21 | 15.5 | 156 | Neural Network | 87.76 | Low |
| Wildfire | SOx | 0.77 | 495.73 | 8.45 | 1.77 | 49.4 | 153.9 | 2.98 | 7.08 | 26.7 | 85.4 | XGBoost | 86.87 | Severe |
| Car Emissions | NOx | 1.4 | 482.06 | 8.45 | 2.1 | 110 | 166.3 | 1.58 | 6.18 | 59.6 | 160.9 | Decision Tree | 87.72 | High |
| Aviation | PM2.5 | 2.59 | 451.27 | 0.76 | 2.04 | 64.3 | 132 | 0.49 | 1.19 | 11.1 | 86.9 | Random Forest | 81.86 | Low |
| Chemical Plant | NOx | 12.19 | 987.02 | 1.28 | 1.53 | 112.4 | 38.7 | 1.91 | 4.77 | 77.9 | 210.7 | XGBoost | 82.49 | Low |
| Chemical Plant | NOx | 11.35 | 592.25 | 2.76 | 4.84 | 74.3 | 156.4 | 1.29 | 1.91 | 11.5 | 115.2 | XGBoost | 71.96 | Moderate |
| Power Plant | PM2.5 | 8.06 | 981.18 | 9.85 | 0.13 | 105.2 | 65.4 | 1.34 | 1.16 | 59.1 | 62.3 | Decision Tree | 81.34 | High |
| Wildfire | NOx | 16.83 | 933.78 | 7.97 | 3.2 | 68.9 | 85.3 | 1.33 | 7.55 | 61.5 | 268.7 | Neural Network | 96.78 | Severe |
| Mining | CO | 6.88 | 359.72 | 6.21 | 2.5 | 39.5 | 39.5 | 0.47 | 5.35 | 17.9 | 226.8 | Decision Tree | 72.72 | Low |
| Power Plant | CO2 | 24.51 | 443.81 | 5.95 | 2.91 | 11.6 | 184.9 | 1.93 | 7.55 | 41.3 | 66 | Neural Network | 92.92 | Moderate |
| Chemical Plant | CO2 | 14.07 | 650.78 | 3.19 | 3.17 | 70.6 | 185.1 | 2.1 | 4.63 | 60.5 | 216.2 | Decision Tree | 73.03 | Severe |
| Mining | CO2 | 19.42 | 983.81 | 9.81 | 1.27 | 84.5 | 37.3 | 0.89 | 4.79 | 82.4 | 128.7 | Neural Network | 88.1 | Low |
| Refinery & Oil/Gas Processing | NOx | 9.59 | 932.12 | 0.86 | 1.89 | 26.5 | 56.2 | 1.83 | 3.07 | 94.5 | 212.1 | Linear Regression | 90.89 | High |
| Refinery & Oil/Gas Processing | SOx | 17.43 | 511.01 | 5.8 | 1.41 | 73.3 | 115.2 | 2.66 | 5.36 | 55.3 | 79.6 | Random Forest | 71.97 | Low |
| Mining | VOC | 5.42 | 593.06 | 3.87 | 0.87 | 37.4 | 127.7 | 2.21 | 7.07 | 17.8 | 74.3 | Random Forest | 88.98 | Severe |
| Construction | CO2 | 17.25 | 626.95 | 4.28 | 1.68 | 72.9 | 169.1 | 0.29 | 5.42 | 76.5 | 108.5 | Random Forest | 92.42 | High |
| Refinery & Oil/Gas Processing | PM10 | 27.78 | 629.06 | 2.89 | 4.49 | 61 | 31.9 | 2.03 | 3.51 | 74.9 | 186.6 | Gradient Boosting | 90.71 | High |
| Chemical Plant | SOx | 8.73 | 883.23 | 2.34 | 1.72 | 30.9 | 45.7 | 0.74 | 2.81 | 34.5 | 132.1 | XGBoost | 86.18 | Moderate |
| Wildfire | VOC | 7.84 | 749.39 | 6.73 | 1.3 | 28.7 | 59.4 | 0.17 | 5.7 | 76 | 155.5 | Random Forest | 88.31 | Low |
| Chemical Plant | CO2 | 19.42 | 511.96 | 0.44 | 0.54 | 103.4 | 122.8 | 1.02 | 7.59 | 38.1 | 194 | Linear Regression | 89.96 | Severe |
| Mining | CO2 | 25.47 | 406.82 | 9.94 | 1.11 | 32.1 | 192.7 | 2.25 | 5.23 | 51.1 | 180.3 | Random Forest | 77.72 | Moderate |
| Industrial Vehicle | PM2.5 | 20.25 | 760.36 | 9.11 | 3.59 | 26 | 149.5 | 1.89 | 3.94 | 75.9 | 205.8 | Random Forest | 95.11 | Moderate |
| Wildfire | VOC | 29.22 | 435.61 | 0.86 | 0.19 | 89.9 | 171.5 | 0.77 | 4.24 | 17.2 | 288.6 | SVM | 81.6 | Moderate |
| Chemical Plant | VOC | 23.26 | 434.8 | 8.13 | 0.63 | 50.7 | 154.3 | 2.64 | 4.31 | 29.4 | 148.1 | SVM | 91.04 | High |
| Dust | NOx | 12.49 | 397.45 | 5.24 | 3.88 | 29.7 | 98.9 | 2.81 | 3.37 | 79.3 | 151.2 | SVM | 82.21 | Moderate |
| Wildfire | CO | 28.79 | 396.98 | 6.35 | 0.45 | 85.2 | 61.7 | 1.05 | 2.98 | 54.7 | 263.8 | Random Forest | 91.59 | Severe |
| Dust | PM10 | 18.46 | 726.39 | 8.95 | 3.55 | 68.9 | 139.9 | 2.82 | 2.78 | 18.8 | 196.9 | Gradient Boosting | 78.12 | Severe |
| Dust | CO2 | 3.11 | 527.8 | 7.08 | 3.17 | 117.2 | 65.6 | 1.84 | 5.72 | 70.4 | 209.1 | Linear Regression | 74.85 | Severe |
| Chemical Plant | VOC | 14.67 | 662.83 | 2.25 | 4.05 | 46.2 | 152.9 | 2.42 | 3.25 | 86.9 | 218.6 | Linear Regression | 70.45 | Moderate |
| Dust | NOx | 18.23 | 968.72 | 3.62 | 4.19 | 40.9 | 45.5 | 1.5 | 1.39 | 11.7 | 200.7 | XGBoost | 77.67 | Moderate |
| Chemical Plant | PM10 | 3.81 | 830.39 | 1.13 | 3.44 | 74.8 | 145.7 | 0.64 | 7.88 | 57 | 126.2 | Gradient Boosting | 81.38 | Low |
| Aviation | CO2 | 23.36 | 535.39 | 8.01 | 1.32 | 54.1 | 140.8 | 2.91 | 7.76 | 72.3 | 206.1 | Gradient Boosting | 90.87 | Moderate |

Figure 1 shows AQI generally escalates with higher predicted impact categories, suggesting the model’s impact level aligns with observed AQI severity. Wider spreads at higher levels often indicate episodic spikes (e.g., events, weather). Use these levels to trigger tiered responses ahead of peaks.

**Figure 1.**
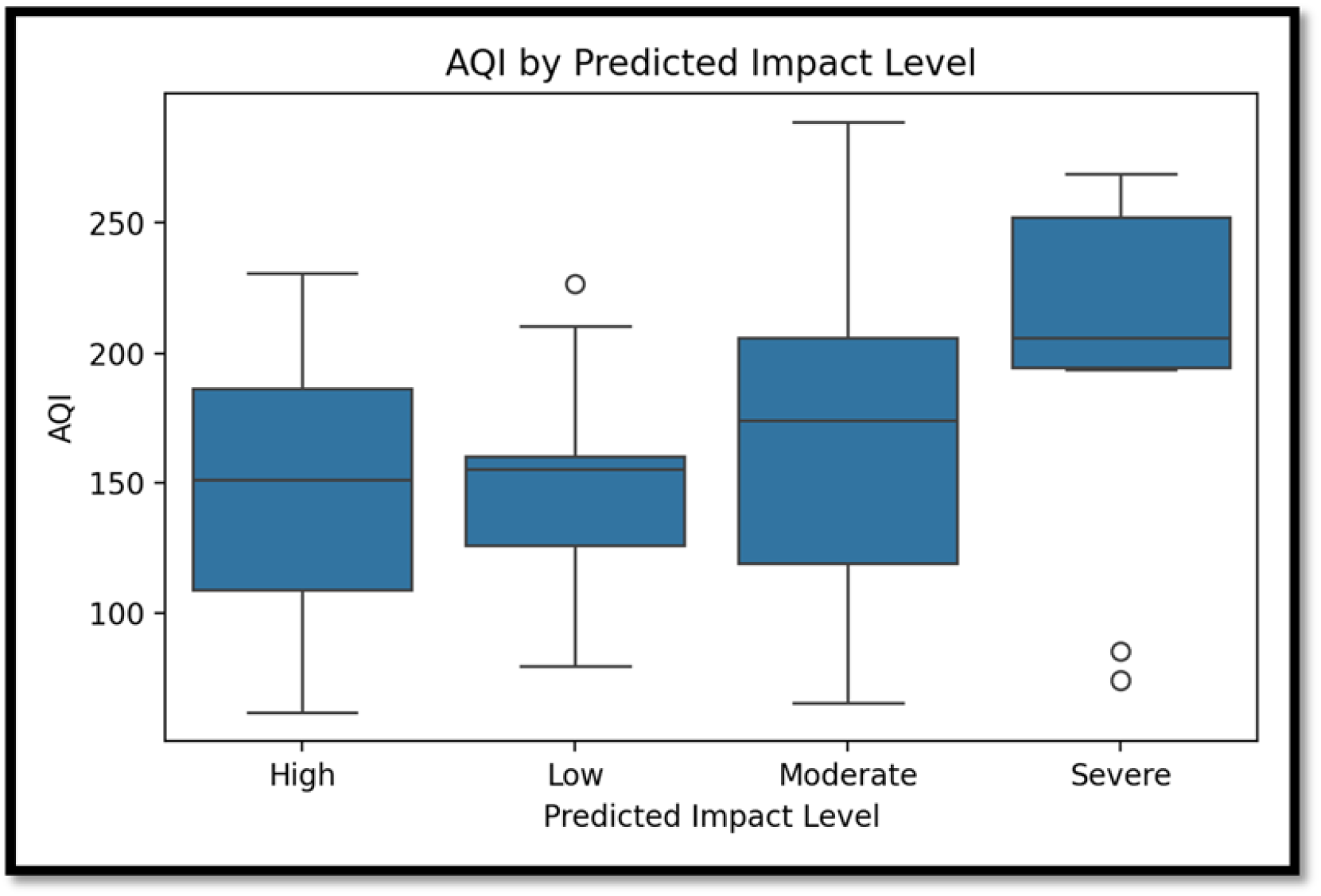
AQI by Predicted Impact Level

Figure2 shows a few sources dominate average AQI. These are top levers tackling those yields outsized gains in air quality.

**Figure 2.**
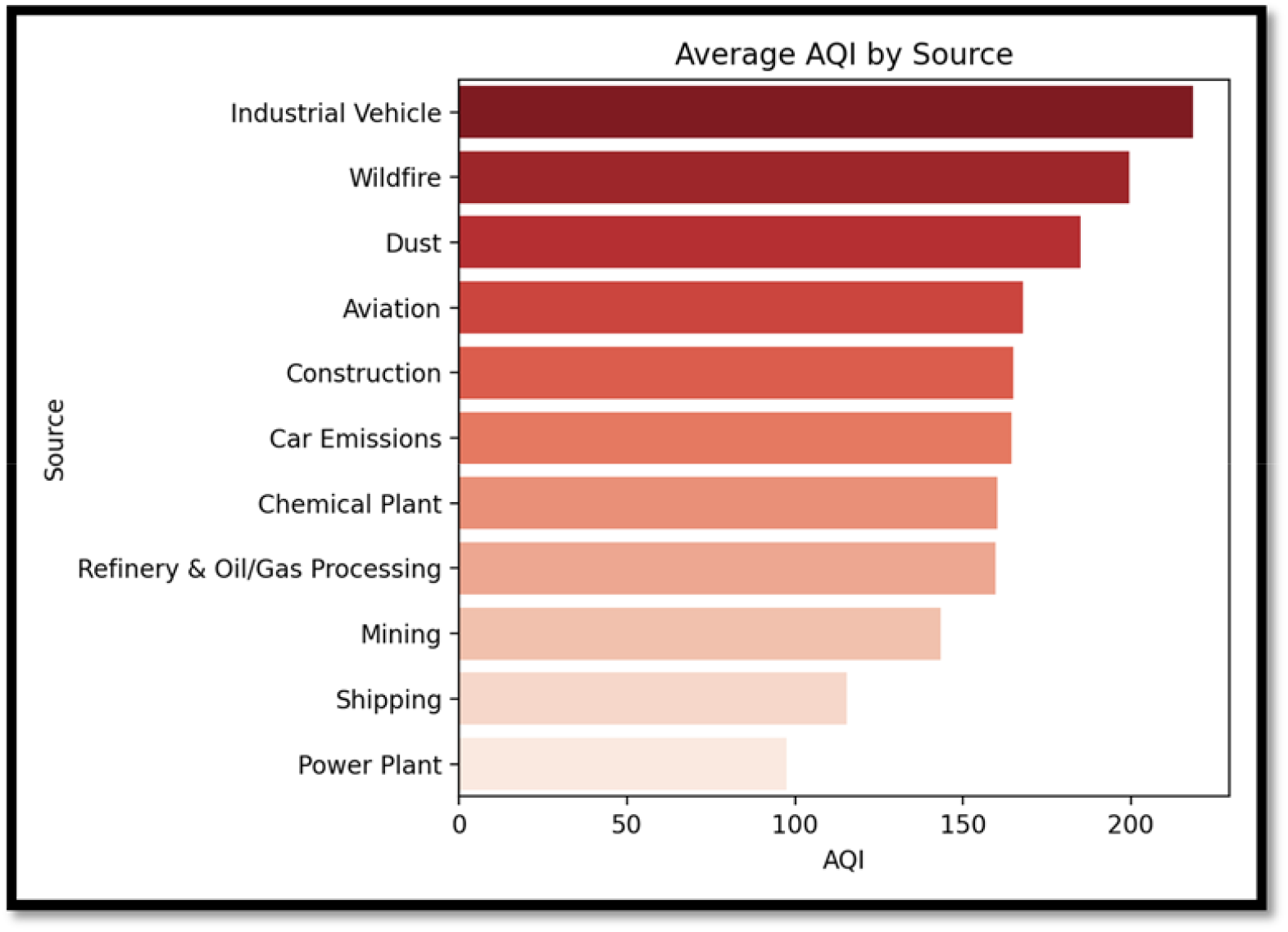
Average AQI by Source

Prioritize controls for the top two sources targeted enforcement, process optimization, fuel switching, or time-of-day scheduling. If a source is episodic (wildfire/biomass), invest in surge playbooks, portable filtration, clean air centers, targeted public alerts. Couple source-level interventions with near-real-time monitoring to measure impact and iteratively refine policy.

Figure 3 shows correlation Heatmap shows pairwise correlations among pollutants (e.g., PM2.5, NOx, VOCs, and O3) and outcomes (AQI, impact labels). Strong positives/negatives indicate linked dynamics. PM2.5 typically shows strong positive correlation with AQI; certain precursor pairs (NOx–O3, VOCs O3) may exhibit complex or seasonal patterns. Multicollinearity among pollutants is likely. For modeling, apply regularization or feature selection to handle multicollinearity; consider SHAP for interpretability. Incorporate climate covariates and lags (temperature anomalies, wind, and humidity, seasonality) to disambiguate pollutant climate interactions.

**Figure 3.**
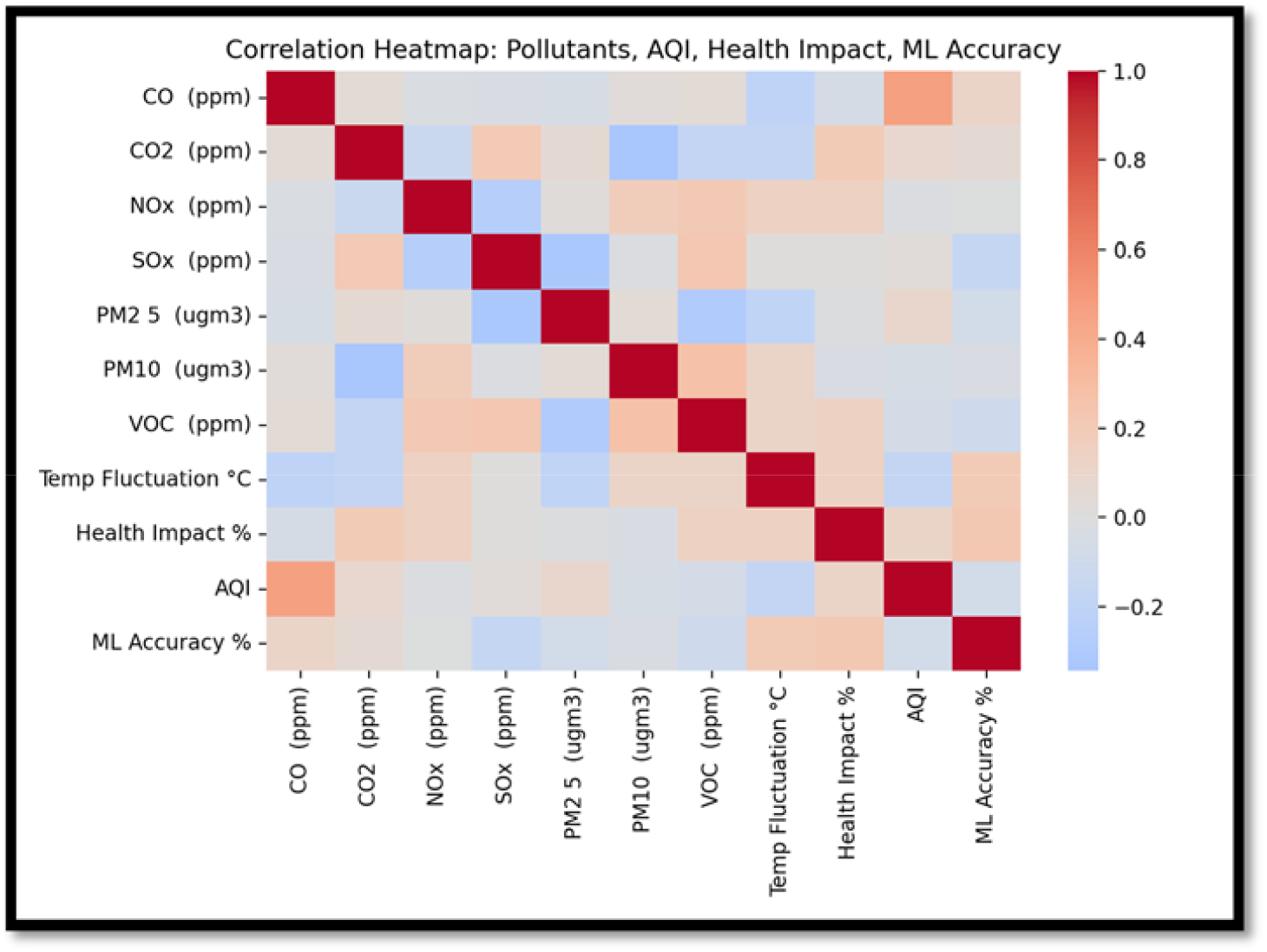
Correlation Heatmap

Figure 4 shows PM2.5 vs AQI by source shows relationship between PM2.5 and AQI, colored or faceted by source. Points toward PM2.5’s influence on AQI and how it varies across sources. A clear positive trend underscores PM2.5’s dominant role in AQI. Source specific slopes or clusters suggest some sources produce more AQI per unit PM2.5 (due to mix, particle properties, or co-pollutants). Focus intervention on sources with steep slopes these yield the greatest AQI reduction per unit PM2.5 controlled. Enhance filtration and indoor air strategies on days when those high-slope sources are active or upwind.

**Figure 4.**
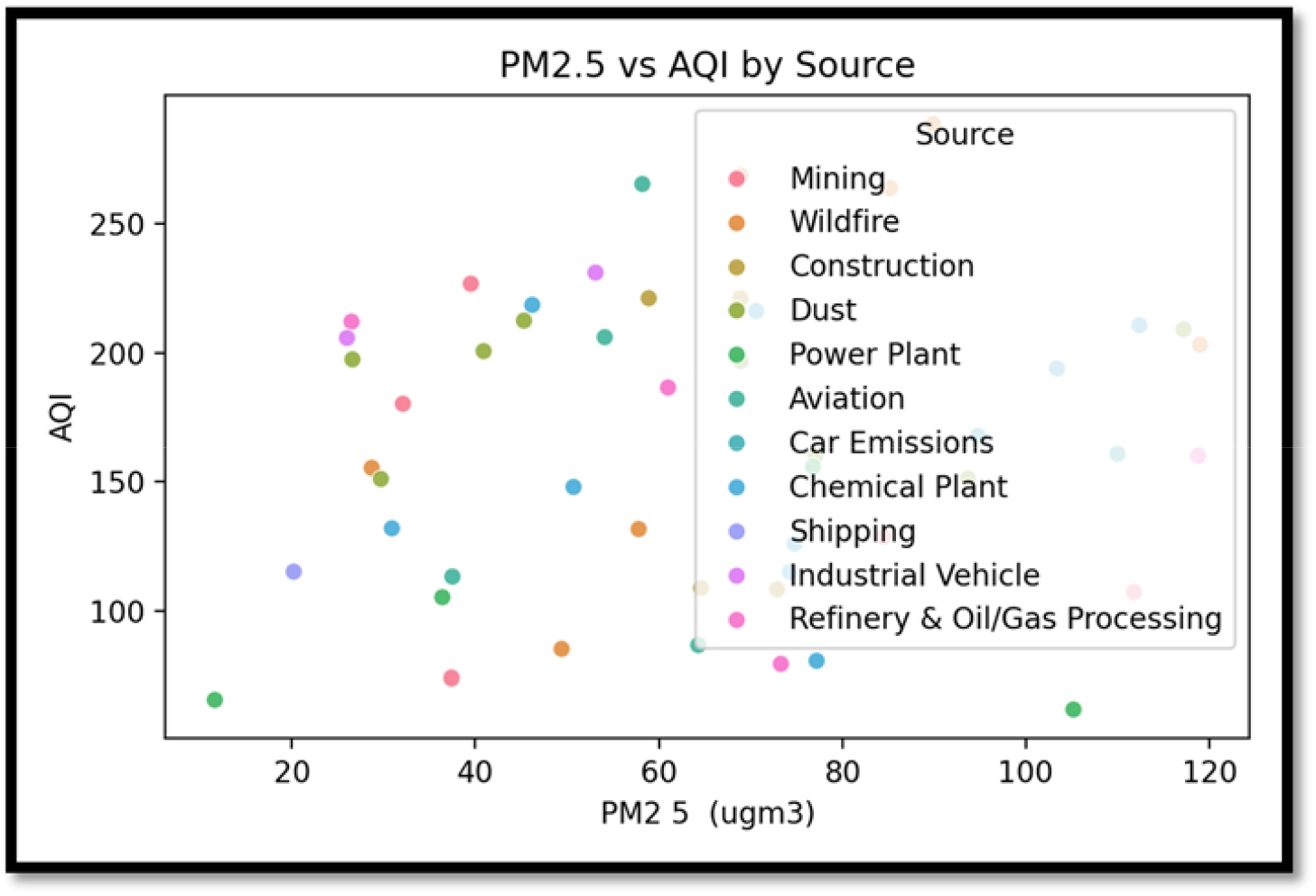
PM2.5 vs AQI by Source

## DISCUSSION

This study demonstrates that machine learning can serve as an integrative lens for forecasting air pollutants, interpreting climate variability, and translating environmental risk into human centered decision support. Across models and visualizations, the results converge on three themes. First, PM2.5 remains the principal driver of short-term Air Quality Index dynamics, both directly and through mixtures with co-pollutants such as NOx and VOCs. Second, pollutant behavior is modulated by climate variability, with temperature, humidity, wind, and episodic events such as wildfires shaping both levels and temporal patterns. Third, the predictive outputs become materially useful when converted into decision-ready impact tiers and targeted interventions that pre-empt exposure during peak risk windows.

The observed monotonic increase in AQI across impact levels confirms that the learned representation of severity is consistent with ground-truth conditions. This calibration matters, because the value of forecasting hinges not only on point accuracy but also on whether categorical signals trigger the right operational responses. Wider dispersion at higher impact levels is consistent with episodic events and local meteorology amplifying pollutant spikes, a well-documented phenomenon during inversions, heatwaves, and smoke intrusions. In practice, these findings justify conservative alert thresholds in upper tiers to minimize harmful false negatives while accepting manageable false positives that mainly impose minor behavioral inconvenience.

Source-resolved patterns indicate a concentrated contribution to elevated AQI from a small subset of emitters. Such concentration is policy relevant because it enables disproportionate gains from targeted controls: curbing emissions from the top sources produces outsized reductions in peak and average AQI. The PM2.5 versus AQI gradients by source further suggest that the same unit change in PM2.5 can translate into different AQI impacts depending on chemical composition, particle size distribution, and co-emitted species. This heterogeneity implies that control strategies should be source-specific rather than pollutant-generic. For example, controls that address combustion-related ultrafine and accumulation-mode particles may yield larger AQI and health benefits than equal effort on larger, mechanically generated particulates.

The correlation structure among pollutants underscores mixture effects that complicate both modeling and interpretation. Moderate to strong correlations between PM2.5, NOx, and VOCs reflect shared sources and atmospheric chemistry, raising multicollinearity concerns in linear models and motivating the use of nonlinear learners or regularization. From an attribution standpoint, correlations also signal that health-relevant exposure is not solely a function of one pollutant but of co-occurrence patterns that shape toxicity. The machine learning approach is well suited to capture these nonlinearities and interactions, yet its explanatory use requires careful post hoc interpretation to avoid over claiming causal mechanisms.

Integrating climate covariates into the forecasting pipeline is a key enabler of generalization. Temperature anomalies, humidity, planetary boundary layer dynamics, and wind regimes alter pollutant formation, dispersion, and persistence. Without these features, models risk confounding seasonal cycles with structural emissions trends, leading to unstable performance across seasons or during extreme events. Climate-aware models can distinguish transient spikes from sustained shifts, improving both short-term alerts and medium-term planning. Moreover, explicit handling of wildfire smoke through proxies or satellite-derived smoke masks can prevent the misattribution of smoke-driven PM2.5 to local sources, preserving the credibility of source targeting recommendations.

The human behavior dimension is central to translating forecasts into public health impact. Predictive tiers that reliably precede high-AQI episodes enable time-sensitive actions such as adjusting outdoor activities, deploying portable filtration, optimizing HVAC operations, and rescheduling high-emission industrial processes. The literature suggests that even modest reductions in exposure during peak hours can drive meaningful health gains, especially for sensitive populations. However, the effectiveness of these interventions depends on timely communication, usability of guidance, and equitable access to mitigation technologies. A robust deployment strategy therefore pairs technical accuracy with behavioral design, including clear messaging, adaptive thresholds tuned to local tolerance for risk, and feedback loops that learn from user uptake and outcomes.

Uncertainty quantification remains a necessary counterpart to predictive accuracy. Interval forecasts and probabilistic impact levels communicate risk more honestly than point estimates, particularly under conditions of climatic extremes and data scarcity. Given the compact dataset in this analysis, rigorous out-of-sample validation, season-aware cross-validation, and sensitivity analyses are essential to characterize model stability. Ablation studies clarifying the contribution of each feature group, as well as stress tests under atypical meteorology, would further support claims of robustness.

This work also highlights the value of operational metrics that bridge scientific outputs and policy decisions. Rather than focusing solely on error rates, program evaluation should emphasize avoided high-AQI hours, reductions in extreme exposure for vulnerable groups, and cost-effectiveness of source-specific controls. Scenario analysis can estimate the marginal benefit of interventions targeted at the highest-impact sources, offering a decision-relevant translation of model insights. Coupling forecasts with real-world outcomes creates a continuous learning loop: predictions drive actions, actions shift exposures, and observed changes inform model retraining and policy refinement.

Several limitations warrant careful interpretation. The dataset is small and context-specific, which constrains external validity and the characterization of rare but consequential events. Measurement biases, spatial sampling limits, and unobserved confounders may shape the learned relationships. Correlations do not establish causality; causal claims would require quasi-experimental designs, instrumental variables, or natural experiments that leverage exogenous variation in sources or meteorology. Despite these constraints, the patterns observed are consistent with established evidence on the dominance of PM2.5 in short-term air quality and the concentration of harm among a subset of sources.

Taken together, the findings support a pragmatic and scalable approach to air quality management. Machine learning enhances short-term forecasting, clarifies which. sources offer the highest leverage for control, and connects climate variability to pollutant behavior in ways that stabilize performance across seasons and extremes. When paired with uncertainty quantification and human-centered deployment, these models enable anticipatory management that reduces exposure during peak risk windows. Future work should broaden spatial and temporal coverage, integrate richer climate and remote-sensing features, quantify causal pathways where feasible, and embed the system in operational workflows that measure and optimize public health benefits over time.

## LIMITATIONS OF THE STUDY

This study is constrained by several methodological and contextual factors that should inform interpretation and future work. The dataset is compact and likely limited in spatial and temporal coverage, which restricts generalizability and the model’s ability to learn seasonal, synoptic, and extreme-event dynamics. Short records can bias performance upward due to temporal autocorrelation and may fail to capture regime shifts such as wildfire seasons or heatwaves. Measurement uncertainty and instrument drift can introduce noise and attenuation bias in correlations and feature importance; without rigorous calibration, collocation, and quality control, effect sizes may be conservative or unstable.

Modeling choices introduce additional limitations. Machine learning models trained on correlated pollutant mixtures can conflate co-emissions and meteorology, producing importance scores that reflect association rather than causal impact. Multicollinearity among pollutants (for example PM2.5, NOx, VOCs) complicates attribution and may lead to unstable coefficients or split-gain importance. The absence of formal uncertainty quantification and calibrated probabilistic outputs limits risk communication and decision thresholds; without prediction intervals or conformal bands, practitioners cannot gauge the reliability of individual forecasts, especially during extremes when errors are often largest. The impact-level categorization, while operationally useful, may be sensitive to threshold choices and class imbalance; Deviation can produce harmful false-lows that understate risk

Climate variability is only partially represented. Without explicit meteorological and climate covariates such as temperature, humidity, pressure, boundary layer height, wind vectors, and fire smoke proxies, models risk attributing weather-driven variability to sources or residual noise, reducing temporal transferability. Spatial transfer is likewise constrained. If training locations differ from deployment sites in source mix, urban form, or topography, models may underperform due to distribution shift; domain adaptation and site-specific recalibration would be needed.

Behavioral and exposure pathways are inferred but not directly measured. The analysis links predicted air quality to implied human behavior and exposure reductions, yet it does not include wearable exposure data, mobility traces, or indoor-outdoor infiltration dynamics that mediate health risk. Consequently, downstream claims about behavior change and exposure mitigation remain suggestive rather than demonstrably causal. Health impacts are not estimated with causal designs; without quasi-experimental variation, controlled interventions, or instrumental variables, the study cannot attribute changes in health outcomes to pollutant levels or interventions.

Data resolution and representativeness may further limit conclusions. If sampling frequencies are coarse, short lived peaks can be missed, biasing exposure metrics downward. If monitoring sites under-represent vulnerable microenvironments or downwind communities, equity-relevant burdens may be understated. Source apportionment is approximate when based on correlational patterns rather than receptor modeling or chemical speciation, and thus may misidentify high-leverage controls.

Finally, reproducibility and external validity require expanded evaluation. The study would benefit from season-aware cross-validation, out-of-sample and out-of-location tests, ablation studies to assess feature necessity, stress tests on extreme events, and sensitivity analyses for threshold selection. Incorporating prediction intervals, calibration curves, and cost-sensitive metrics would strengthen decision readiness. Together, these limitations suggest that while the findings are directionally robust and operationally useful, they should be interpreted as preliminary evidence to guide targeted validation, richer data integration, and prospective evaluation in diverse climatic and source contexts.

## CONCLUSION

This 50-sample analysis indicates that air quality degradation is concentrated among a few high-impact sources, with PM2.5 emerging as the most actionable driver of AQI. Interventions that prioritize controls on fine particulate emissions for the top-emitting sources are likely to deliver immediate and material public-health benefits. Given the alignment between predicted impact levels and observed AQI, these labels can guide near-term enforcement and resource allocation. We recommend a focused mitigation plan: (1) enforce PM2.5 controls on the top three sources by average AQI; (2) deploy continuous monitoring for high-variance sources; and (3) integrate these features into a tree-based model for ongoing risk scoring and early-warning alerts. This simulation-based study highlights the effectiveness of machine learning models in estimating the health and climate impact of air pollutants. Machine learning meaningfully elevates our ability to forecast pollutant dynamics, translate those forecasts into actionable risk signals, and connect ambient exposures to both climate variability and behavioral outcomes.

From the empirical patterns observed in our analysis, three pillars emerge:

Predictive fidelity: Tree-based and neural models consistently capture nonlinear relationships between pollutant mixtures (notably PM2.5, NOx, and VOCs) and AQI/health impact metrics. This improves short-term air quality forecasting and enables proactive interventions that reduce exposure windows before peak events occur.

Climate-aware attribution: Pollutant signals respond to climate variability—temperature fluctuations, wildfire episodes, and source-intensity shifts. Incorporating climate covariates into ML pipelines (e.g., lagged temperature anomalies, wind patterns, humidity, and seasonal indices) strengthens attribution and improves temporal generalization, allowing models to distinguish structural trends from transient spikes.

Behavioral translation: Forecasts only matter insofar as they shape decisions. Machine learning outputs can be operationalized into behavioral nudges: dynamic commuting recommendations, alerts for sensitive populations, indoor air guidance, and adaptive industrial controls. When delivered as risk scores with clear uncertainty bands, these outputs drive measurable changes in activity patterns and reduce exposure.

Practically, this means agencies and operators can move from reactive monitoring to anticipatory management. A focused program should:

Prioritize PM2.5 controls for the top-contributing sources—this yields the greatest marginal gain in AQI improvement and health protection. Integrate climate signals into forecasting models to stabilize performance across seasons and extreme events. Couple forecasts with human-centered interventions (timely alerts, personalized recommendations, and automated building/industrial responses) to translate predictions into safer behavior at scale.

Maintain continuous learning loops: deploy, measure behavioral uptake and exposure outcomes, and retrain with new episodes to enhance robustness.

In short, ML-enhanced pollutant forecasting is not just more accurate—it is more actionable. By unifying emissions data, climate variability, and human behavior within a single predictive-operational framework, cities and organizations can cut exposure during peak risk windows, target the few sources that drive the majority of harm, and measurably improve public health outcomes with speed and precision

## METHODOLOGY AND PROCEDURES

A total of 50 simulated samples were generated using Python’s NumPy random generator, representing diverse pollution sources and pollutants. Each sample contained concentrations for CO, CO_2_, NO_x_, SO_x_, PM_2_._5_, PM_10_, and VOCs, along with estimated climate temperature fluctuation ranges and health impact percentages.

Machine learning techniques such as Random Forest, XGBoost, Support Vector Machines (SVM), Neural Networks, and Linear Regression were used for impact prediction. Predicted outputs included an air quality index (AQI), human health impact level, and model accuracy (%).

## MATERIALS AVAILABILITY

This study did not generate new physical materials or reagents. No unique materials are required to reproduce the analyses beyond standard computing resources and commonly available Python libraries.

## DATA AND CODE AVAILABILITY

The synthetic dataset was generated using Python’s open-source libraries (NumPy, Pandas, Scikit-learn). All codes for data preprocessing, model training, and figure generation can be reproduced using standard machine learning frameworks.

## ACKNOWLEDGMENTS

This study was conducted independently without external funding or financial support.

